# Mining drug-target interactions from biomedical literature using chemical and gene descriptions-based ensemble transformer model

**DOI:** 10.1101/2023.07.24.550359

**Authors:** Jehad Aldahdooh, Ziaurrehman Tanoli, Jing Tang

## Abstract

Drug-target interactions (DTIs) play a pivotal role in drug discovery, as it aims to identify potential drug targets and elucidate their mechanism of action. In recent years, the application of natural language processing (NLP), particularly when combined with pre-trained language models, has gained considerable momentum in the biomedical domain, with the potential to mine vast amounts of texts to facilitate the efficient extraction of DTIs from the literature. In this article, we approach the task of DTIs as an entity-relationship extraction problem, utilizing different pretrained transformer language models, such as BERT, to extract DTIs. Our results indicate that an ensemble approach, by combining gene descriptions from the Entrez Gene database with chemical descriptions from the Comparative Toxicogenomics Database (CTD), is critical for achieving optimal performance. The proposed model achieves an F1 score of **80.6** on the hidden DrugProt test set, which is the top-ranked performance among all the submitted models in the official evaluation. Furthermore, we conduct a comparative analysis to evaluate the effectiveness of various gene textual descriptions sourced from Entrez Gene and UniProt databases to gain insights into their impact on the performance. Our findings highlight the potential of NLP-based text mining using gene and chemical descriptions to improve drug-target extraction tasks. Datasets utilized in this study are accessible at https://dtis.drugtargetcommons.org/.

## Introduction

The study of small molecules and their interactions with protein targets is a key element in drug discovery research. Over the years, a vast amount of information about these interactions has been published in scientific literature including high throughput screening studies [1, 2]. With recent advancements in NLP technologies, there is an ever-increasing need to develop semi-automated text-based transformer models (such as BERT) that can dig further into the text of a screening study to mine DTIs subject to human approval [3]. In the field of NLP, entity relationship extraction is a process of identifying and extracting meaningful relationships between entities (in this case compounds and protein targets) from textual data. To accelerate the extraction of drug-target interactions (DTIs), the BioCreAtIvE (Critical Assessment of Information Extraction systems in Biology) consortium organized a DrugProt challenge [4], which provided participants with a manually annotated dataset, the DrugProt corpus, which contains PubMed abstracts, manually annotated chemical compound and gene/protein mentions, and their interactions of 13 types that cover key relations of biomedical importance.

Most of the participants in the DrugProt challenge applied the most recent pre-trained models like BERT to train on the domain of drug-target interactions. For example, in [5], authors leveraged ensembles of pre-trained language models such as RoBERTa-large [6], using chemical descriptions to achieve an F1 score of 79.73 as the best model in the DrugProt challenge. More recently, authors in [7] employed a sequence labelling framework, achieving a state-of-the-art F1-score of 80.0 on the test set.

In a recent study [8], the authors proposed to enhance the supervised extraction of DTIs by integrating distantly supervised models. While traditional supervised methods rely on manually labelled data for training, which can be resource-intensive and time-consuming, distantly supervised models utilize existing knowledge bases to label the data automatically. However, their experimental results revealed that the proposed approach could have improved the performance accuracy as expected. On the contrary, the inclusion of mislabeled data resulted in even lower performance accuracy. Therefore, this study highlights the limitations within the current supervised approach to DTIs extraction.

In light of the challenges faced by the supervised extraction approaches, an alternative strategy in [9] has emerged which emphasizes the integration of heterogeneous knowledge graphs (HKGs) into drug-drug interaction extraction from biomedical literature. Recognizing the necessity of incorporating a diversity of background knowledge into the extraction process, like human experts comprehend and dissect the scientific literature, the authors built the drug representations by conducting a link prediction task on a pharmaceutical HKG dataset. Then, they combined the contextual information of the input sentences in the corpus with the drug information from the HKG dataset. On evaluation with the DrugProt dataset, their method achieved an F1-score of 77.9 on the development set.

In this study, we delve into a distinctive aspect of gene descriptions sourced from the Entrez Gene database—an avenue not previously explored, as most research primarily focuses on utilizing information from protein databases such as UniProt [10]. Additionally, we pioneered the combined use of both chemical and gene descriptions. To achieve this, we deployed a Multi-Model Ensemble (CGDE) that combines specialized models focused on analyzing chemical and gene data. Our work not only improves the F1 scores achieved by the aforementioned related works but also highlights the importance of incorporating gene descriptions in the DTI extraction process. By leveraging the rich information in gene descriptions, our method demonstrates a more comprehensive understanding of the intricate relationships between drugs and their target proteins, ultimately contributing to more efficient and accurate DTI extraction from biomedical text.

## Materials and methods

### Datasets

We have mainly used DrugProt for training the models. DrugProt is a publicly available dataset containing manually annotated entities and relations relevant to drug discovery research. The DrugProt dataset consists of training, development, and testing subsets. Although the actual labels for the test dataset remain hidden, participants can submit their predictions to the Codelab platform and receive the final prediction results. Figure 1 provides a comprehensive overview of the statistics for the DrugProt training and development datasets. Within this dataset, gene entities are categorized into two primary classifications: GENE-Y, which represents normalizable genes, and GENE-N, marking non-normalizable genes. In terms of relation types, the DrugProt dataset is quite extensive, featuring 13 distinct categories, including: INDIRECT-DOWNREGULATOR, INDIRECT-UPREGULATOR, DIRECT-REGULATOR, ACTIVATOR, INHIBITOR, AGONIST, ANTAGONIST, AGONIST-ACTIVATOR, AGONIST-INHIBITOR, PRODUCT-OF, SUBSTRATE, SUBSTRATE_PRODUCT-OF, and PART-OF. These classifications enable a comprehensive understanding of the relations and interactions among various entities within the DrugProt dataset.

**Figure 1.**
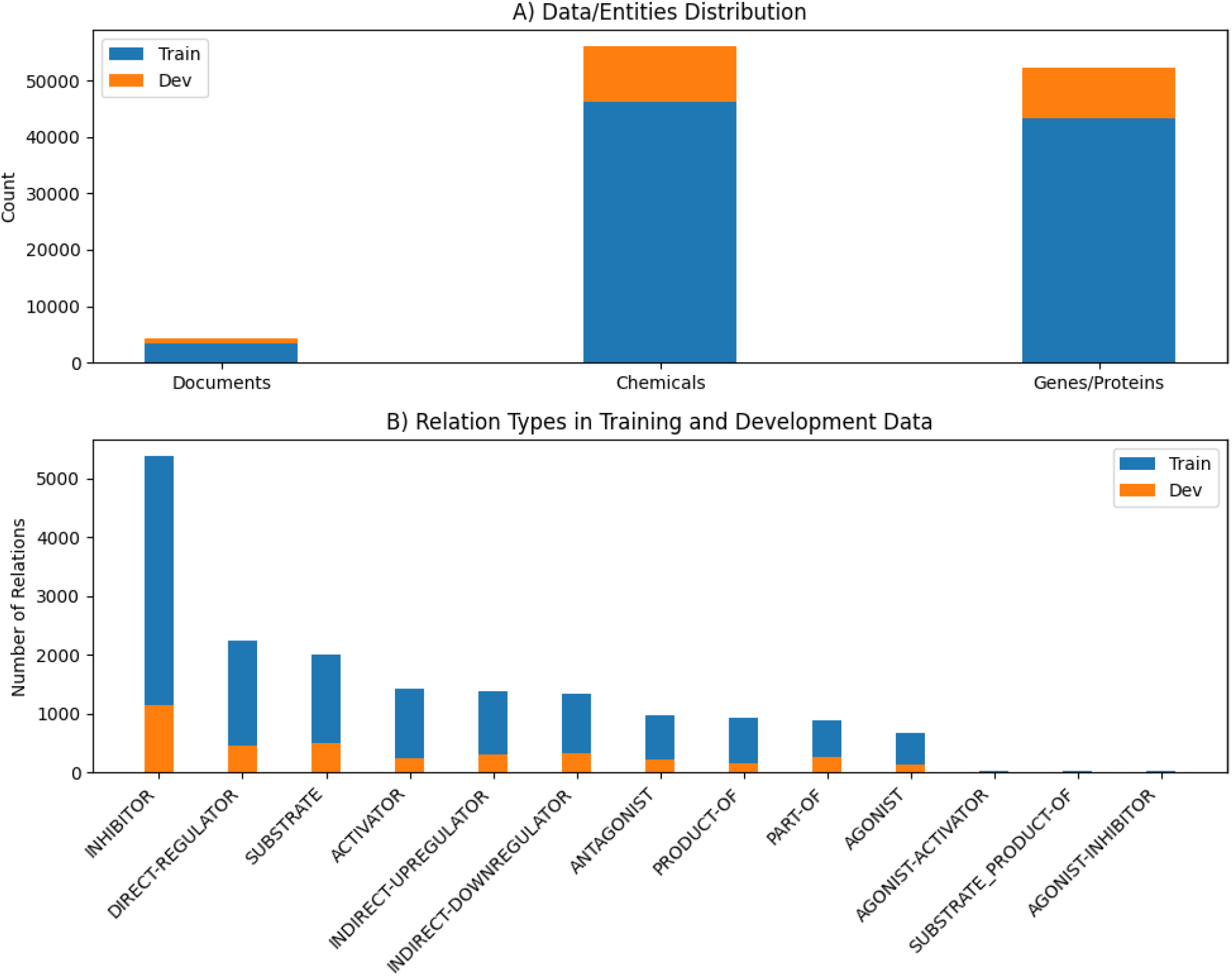
Distribution of the biological entities in the DrugProt dataset. (A) Number of documents (PubMed articles), chemical and gene entities. (B) Number of relation types in DTIs.

We conducted experiments using various model configurations, and subsequently linked chemical and gene to their ontologies. To ensure consistency, we adopted the same unique gene identifiers as in [5] for both the training and development datasets. These identifiers were normalized using a BioSync model [25] for chemicals, leveraging the comprehensive BioCreative V CDR (BC5CDR) dataset [26], and for proteins, utilizing the BioCreative II Gene Normalization (BC2GN) dataset [27].

To retrieve chemical and gene entities for the hidden test dataset, we utilized the Pubtator’s API [11], which utilizes GNormPlus [12] for the gene annotation and determines their NCBI Gene identifiers (Figure 2). For the chemical annotation, Pubtator employs TaggerOne [13] to link the chemical names to MeSH identifiers. We have also employed the BioSync method [25] to perform the normalization, using dmis-lab/biosyn-sapbert-bc2gn model, which was applied to the BC2GN dataset. Using the gene IDs, we obtained the gene summary descriptions from the Entrez Gene database, making our experiment the first attempt to acquire such information. Furthermore, we extracted the description for each gene from their respective web pages using specific HTML tags. For the protein annotation, we obtained the descriptions from the UniProt database.

**Figure 2.**
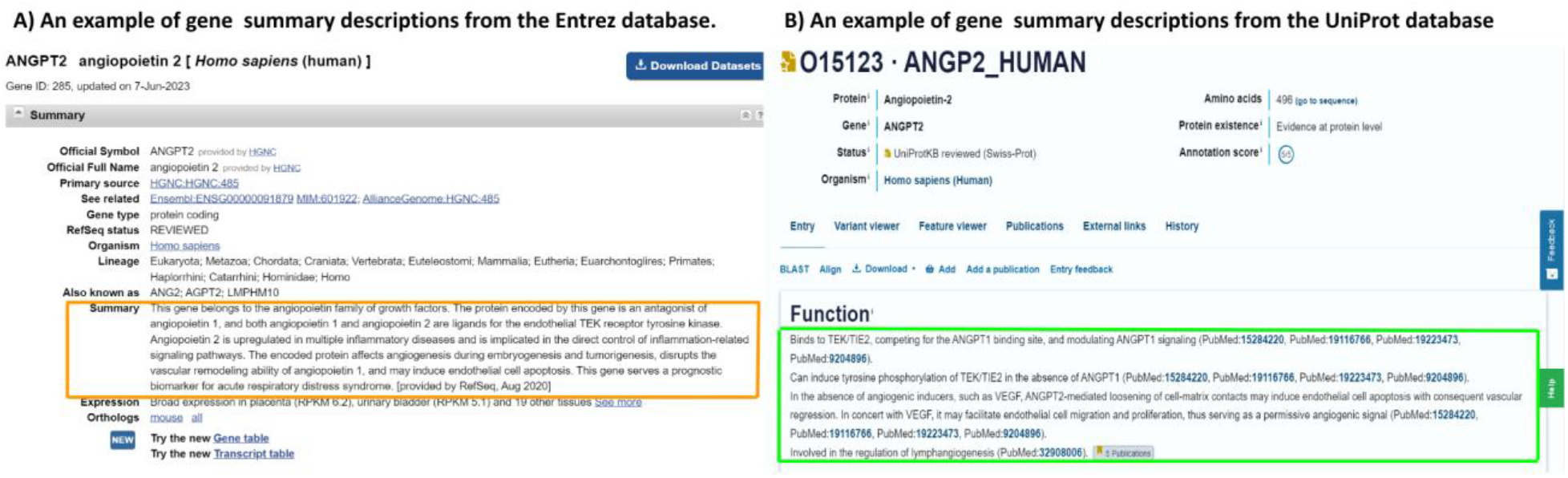
An example of gene/protein summary and function descriptions from the Entrez (A) and UniProt (B) databases. Extracted text descriptions from both databases are highlighted in the rectangle.

In our study, we utilized the same chemical descriptions as in [5], which were derived from the Comparative Toxicogenomics Database (CTD). These descriptions were obtained from the Definition field of the CTD’s chemical vocabulary, specifically taking the first sentence of the definition. Furthermore, we computed **576** descriptors from the gene sequences using Amino Acid index database [14]. From these, 420 are based on monopeptide and dipeptide frequencies for the 20 amino acids. The remaining 156 features relate to various physicochemical properties, mass, length and free energy. To facilitate a comprehensive understanding and replication of our computational methodology, we have developed a Python script that automates the extraction and calculation of these descriptors. The script employs the Biopython library [28] to analyze protein sequences, calculating a broad array of features such as molecular weight, aromaticity, hydrophobicity, flexibility, polarizability, free energy, and steric properties, among others. It includes functions for counting amino acids, computing their percentages in sequences, and assessing protein stability through the instability index, gravy (grand average of hydropathicity), and isoelectric point calculations. This script, accessible via our GitHub repository (https://github.com/JehadAldahdooh/BioSeqFeatureExtraction). By detailing the computational steps and parameters used to derive our dataset’s descriptors, we aim to enhance the reproducibility and transparency of our research methodology.

## Methods

We approached the drug-target relation classification task as a multi-label problem similar to [5]. However, we employed various setup configurations, including the use of Focal Loss instead of binary cross-entropy loss. Furthermore, we incorporated both gene and chemical description datasets. We utilized various pre-trained BERT (Bidirectional Encoder Representations from Transformers) models (large and base), including SciBERT [15], BioBERT [16], RoBERTa [6], BioLinkedBert [17], with a primary emphasis on the RoBERTa model. We combined all these models to form an ensemble of different transformer models.

SciBERT [15] is a scientific text-specific version of BERT, fine-tuned on an expansive corpus of scientific papers from Semantic Scholar, encompassing diverse scientific disciplines. SciBERT is designed to better capture and understand the language and terminology used in scientific literature. In our experiments, we employed the allenai/scibert_scivocab_uncased (https://huggingface.co/allenai/scibert_scivocab_uncased) model, which is publicly available through the Hugging Face Hub (https://huggingface.co/models). BioBERT [16] offers a specialized language representation model for the biomedical domain, trained on large-scale biomedical corpora, available at dmis-lab/biobert-large-cased-v1.1 (https://huggingface.co/dmis-lab/biobert-large-cased-v1.1) model. RoBERTa (Robustly optimized BERT) is a variant of the BERT model developed by Facebook AI and is designed to improve upon the original BERT model in several ways by using a larger and more diverse training datasets, longer training time, and dynamic masking to improve the model’s ability to understand natural languages. RoBERTa also removes the next sentence prediction objective used in BERT, which was found to be less important for many NLP tasks. RoBERTa-large is a variant of the RoBERTa model, with even more parameters and a longer training time. It is designed to further improve upon the original RoBERTa model’s ability to understand natural languages. RoBERTa-large is generally considered to be more powerful and accurate than the smaller RoBERTa model. We used the RoBERTa-large-PM-M3-Voc [18] (https://github.com/facebookresearch/bio-lm) model in our experiments.

BioLinkBERT [17] is a unique version of BERT that includes document link knowledge, such as hyperlinks and citation links from PubMed abstracts, allowing the model to integrate information from multiple documents. Pre-training involves feeding linked documents into the same language model context, in addition to a single document. In our experiments, we employed the BioLinkBERT-large model that is available at (https://huggingface.co/michiyasunaga/BioLinkBERT-large).

To generate the training and development instances from an abstract, we considered all compound-target pairs that occur in the same sentences. We created one instance for each compound-target entity pair, inserting special tokens [HEAD-S], [HEAD-E], [TAIL-S], and [TAIL-E] to mark the start and end of the head (the chemical entity) and tail (the target gene entity). We used the context embeddings of the [CLS] token from the last hidden layer of the BERT model and fed it to a fully connected layer to obtain the logits. Additionally, we experimented with the embeddings of the target gene features, concatenating them with the [CLS] output. The main framework is shown in Figure 3. Moreover, we incorporated an experimental strategy of varying random seeds to assess their impact on the model’s stability and performance. The random seeds used were: 0, 1, 3, 4, 5, 7, 9, 10, 20, 42, and 1024.

**Figure 3.**
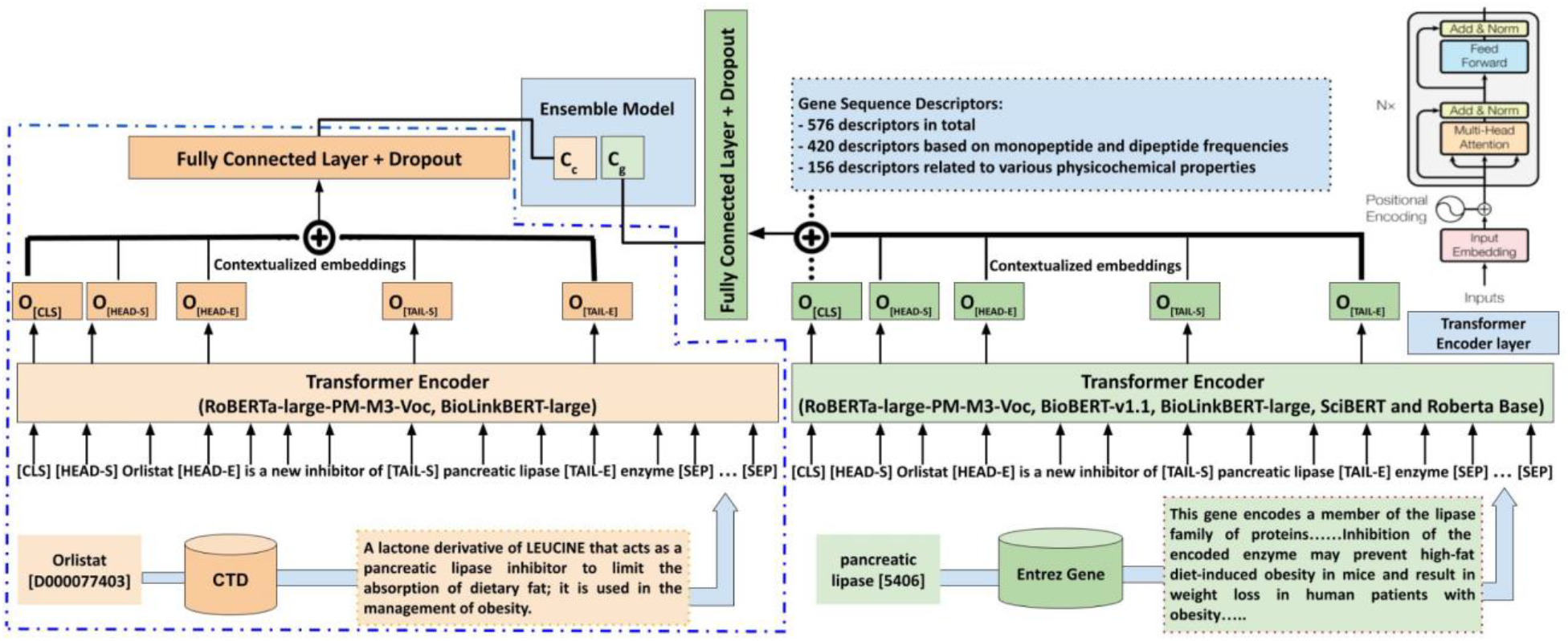
Overview of the best performing ensemble model. Our experiments utilize an ensemble of the RoBERTa-large-PM-M3-Voc, BioBERT-v1.1, BioLinkBERT-large, SciBERT, and Roberta Base models. The abstract texts are processed with BERT, together with target gene and chemical descriptions extracted from the Entrez and CTD databases, respectively. Furthermore, we integrated 576 gene sequence-based features. Special tokens mark the start and end of chemical-gene entities. These tokens are also incorporated in the model’s input. The left side of the diagram is represented by blue dashed lines, which emphasize the model’s focus on chemical descriptions, whereas the right-side section is dedicated to utilizing gene descriptions and features based on gene sequences.

### Loss function

We used Focal Loss, which is a loss function designed to address the issue of class imbalance in classification tasks. Focal Loss was first introduced in [19] to improve the performance of object detection models. However, it can also be applied to our classification problem where there is a class imbalance between the relations (Figure 1). The Focal Loss is an extension of the Binary Cross Entropy (BCE) loss, but it introduces a modulating factor that decreases the contribution of well-classified examples to the loss. This helps the model to focus more on the hard-to-classify examples, thus addressing the class imbalance issue. Focal loss has two main parameters:

- Gamma (γ): The focusing parameter, which determines how much the contribution of well-classified examples is reduced. A higher value of gamma will lead to a more focused loss function. In our study, we set the focusing parameter to γ = 0.5.
- Eps (ε): A small value added to the logarithm term to prevent division by zero or taking the logarithm of zero. We used the default value 1e-6.

It is defined as:

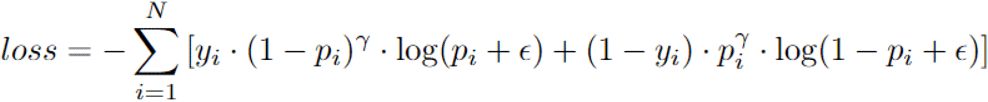

where:

- N is the total number of samples in the dataset
- y_i is the true label for the ith sample
- p_i is the predicted probability for the ith sample

### Experiments

We ran the experiments with 10 different random seeds and averaged the results to obtain a more robust estimate of the prediction performance. To assess the effectiveness of our approach, we conducted a series of four experiments. Subsequently, we submitted the final predictions to the official site for the evaluation on the hidden test set. The results were reported as micro-averaged F1 computed using the official DrugProt evaluation library (https://github.com/tonifuc3m/drugprot-evaluation-library). The experiments conducted are summarized as follows:

#### Model 1 (Base model)

Sentence-Based Ensemble (SBE) Model

In this experiment, we trained an ensemble model consisting of 10 RoBERTa-large-PM-M3-Voc models, with different seeds. We exclusively utilized sentences without any external chemical or gene descriptions.

#### Model 2

Gene Description Ensemble (GDE) Model

The main goal of this experiment was to assess how gene descriptions affect prediction accuracy. To achieve this, we constructed a Gene Description Ensemble (GDE) comprising 10 models. These models encompassed different models to leverage their unique strengths – consisting of the Roberta Base, SciBERT, RoBERTa-large-PM-M3-Voc, BioLinkBERT-large, and BioBERT-v1.1 single models – all of which incorporated gene descriptions. By including a diverse range of model architectures and learning strategies within the GDE, we aimed to provide extensive coverage and enhance the ensemble’s overall performance. The ensemble model is constructed by taking the average of the probabilities generated by the models with a threshold of 0.5 or higher.

#### Model 3

Chemical Description Ensemble (CDE) Model

To establish the benchmark for our comparative analysis, we replicated the top-performing model from the DrugProt Challenge, which achieved an F1 score of **79.7** on the concealed DrugProt test data. This result was attained by training an ensemble of ten RoBERTa-large-PM-M3-Voc models, incorporating chemical definitions from the Comparative Toxicogenomics Database (CTD) for both the training and development sets.

We employed the same approach by developing an ensemble of 10 RoBERTa-large-PM-M3-Voc models with different seeds. The ensemble model we employed – consisting of the RoBERTa-large-PM-M3-Voc and BioLinkBERT-large single models.

#### Model 4

Chemical and Gene Description Ensemble (CGDE) Model

We employed an ensemble that utilized both gene and chemical descriptions models as an integration to models 1 and 2.

#### Model 5

Gene sequence-based Features and text Descriptors Ensemble (GFDE) Model

Gene sequence-based features were incorporated in this experiment. An ensemble of 10 RoBERTa-large-PM-M3-Voc models with different seeds has been constructed using gene descriptions, along with gene sequence-based features. Nevertheless, they did not contribute to an improvement in the results. In order to preprocess the gene sequence-based features, L2 normalization was applied by dividing each vector by its L2 norm, as expressed in Equation (2).

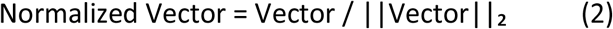

Where:

- Normalized Vector represents the resulting vector after applying the normalization process.
- Vector represents the original vector.
- ||Vector||_2_ represents the L2 norm of the vector.

## Results and discussion

### Performance of the five different models on the hidden DrugProt test data

The five models demonstrated varying levels of performance on the DrugProt hidden test data. The CGDE model achieved the highest F1 score of **80.6**, followed by the GDE model with 80.1, the CDE model with 79.7, and the GFDE model with 79.5. Additionally, the SBE model exhibited an F1 Score of 77. These results indicate that the CGDE model outperformed the other models in terms of overall predictive accuracy (Figure 4). Our experiments showed that utilizing only the Entrez gene descriptions (GDE Model) yielded better performance, with an improvement by approximately 0.4 F1 compared to the CDE baseline model. Combining chemical and gene descriptions led to a more significant increase, with approximately 0.9 points F1 improvement over the CDE baseline model. These results suggested that integration of chemical and gene descriptions in an ensemble BERT model achieved the best performance, compared to the top-performing model from the DrugProt Challenge. However, adding gene sequence-based features besides gene descriptions led to a slight decrease of approximately 0.2 points F1, suggesting that the numeric features may not be optimal when combined with BERT contextual embeddings. In Figure 5, we compared the performance per class achieved by the five different models.

**Figure 4.**
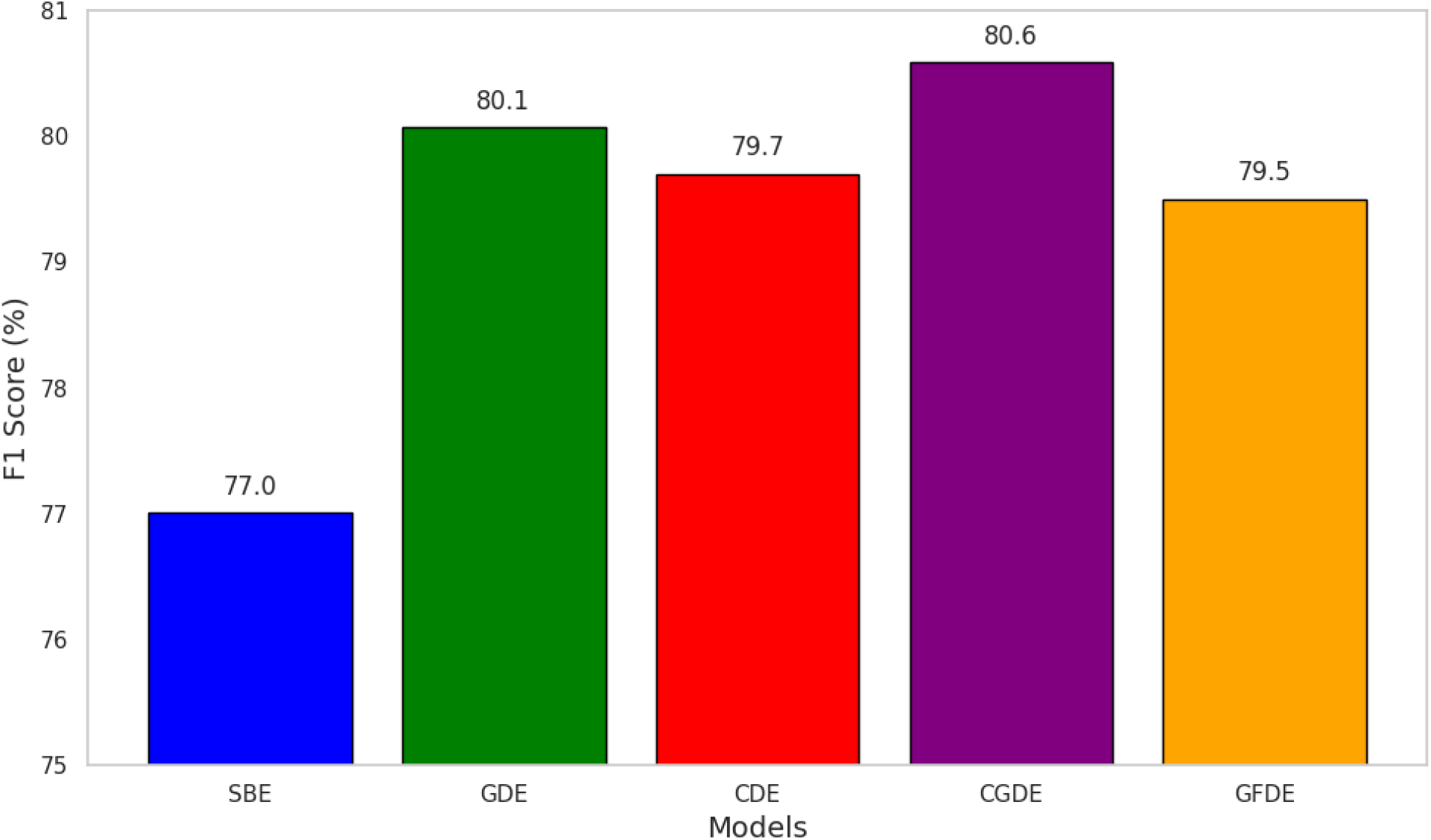
Performance comparison of the five models on the hidden DrugProt test set, presented as micro-averaged F1 scores.

**Figure 5.**
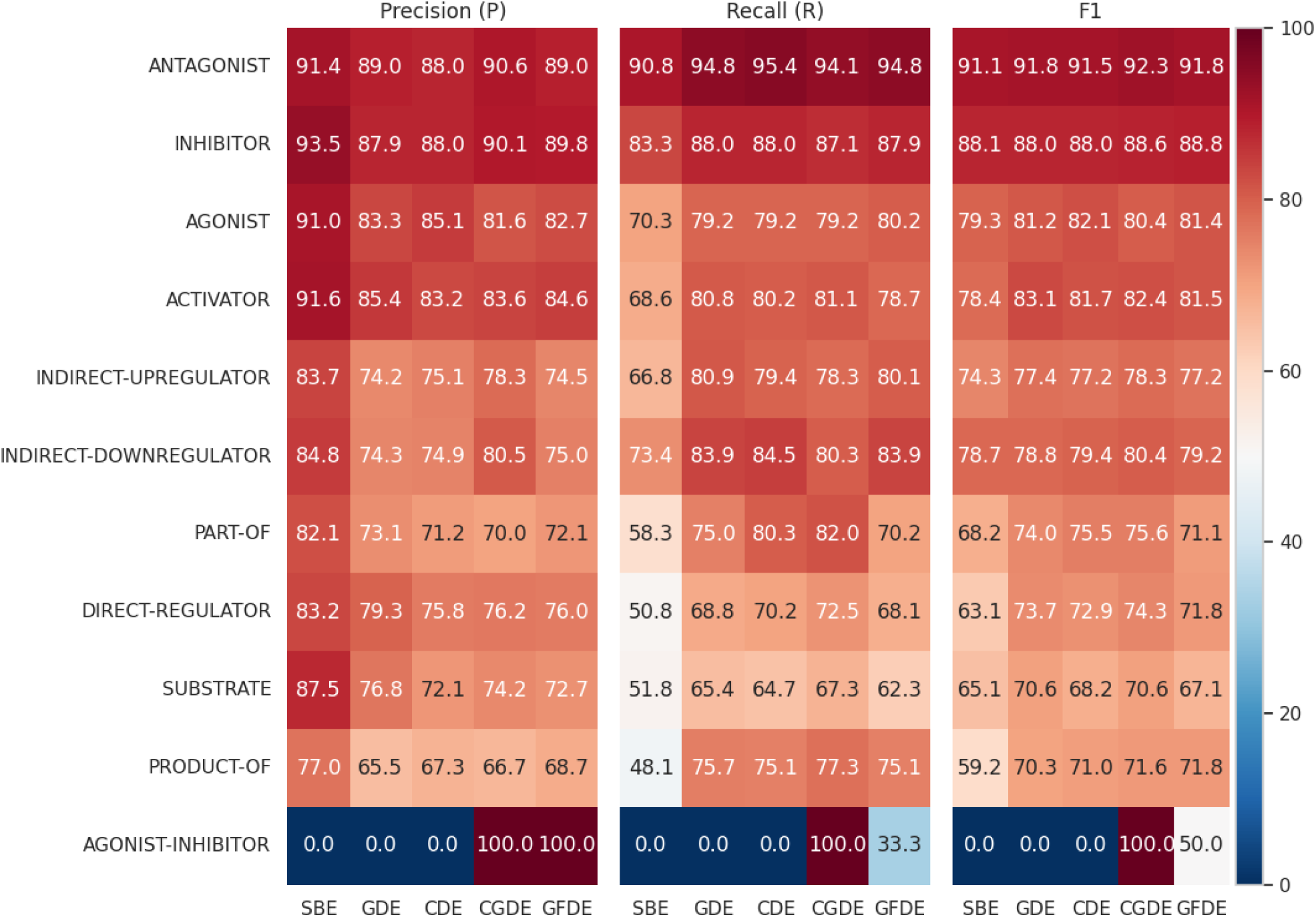
Comparative analysis of model performance per relation types.

It can be shown that the ANTAGONIST relation seems to consistently have the highest F1 score across all models, indicating that this class was the easiest for the models to predict correctly. The highest F1 score is 92.3 by the CGDE model. Furthermore, both the ANTAGONIST and INHIBITOR interaction types display a stable and consistent performance across all the five models. This could suggest that the features for these types of interactions are robust and well-captured, which consistently rank in the top 2 in terms of precision, recall and F1 scores across all the models. The CGDE model generally performs the best across different interaction types. It achieves the highest F1 score for ANTAGONIST, DIRECT-REGULATOR, SUBSTRATE, PART-OF, INDIRECT-UPREGULATOR, INDIRECT-DOWNREGULATOR, and AGONIST-INHIBITOR. However, the GFDE model performs better for INHIBITOR, and PRODUCT-OF relations.

#### Performance on rare classes

The AGONIST-INHIBITOR relation shows the most variability in accuracy across different models since it is rarely present (Figure 1). For example, the GFDE model has an F1 score of 50, CGDE has an F1 score of 100, while SBE, GDE and CDE have an F1 score of 0. This discrepancy could be due to the different ways these models handle this specific interaction class.

In the case of CGDE, we implemented a lower threshold (>= 0.2 instead of >= 0.5) to detect this relation. This adjustment was necessary due to its relatively low representation in the training and development dataset, as well as the model’s tendency to predict this relation with a lower probability. Conversely, in the GFDE model, the gene features appear to aid in capturing the context of the agonist-inhibitor relation, allowing the model to identify it without altering the threshold.

Additionally, none of the five models predict AGONIST-ACTIVATOR or SUBSTRATE_PRODUCT-OF relations, primarily due to the limited size of training and development samples available for these specific relationships (Figure 1).

#### Recall and Precision trade-off

There is an evident trade-off between precision and recall in the five different models. For example, in the DIRECT-REGULATOR relation, the SBE model has a high precision of 83.2 but a relatively low recall of 50.8. In contrast, the CGDE model has a lower precision of 76.2 but a higher recall of 72.5. This indicates that SBE is more conservative in predicting DIRECT-REGULATOR, resulting in fewer false positives but more false negatives. The CGDE, on the other hand, makes more positive predictions, leading to fewer false negatives but more false positives.

### Performance comparison with the latest top-performing model on DrugProt

After the DrugProt Challenge, more models have been submitted, with one of them outperforming the original top-performing CDE model. The latest top-performing model was achieved by the NLM-NCBI team, who utilized an ensemble approach employing sequence labeling models with standard loss that were trained using DevES and TrainES via majority voting [5]. Based on the results of the hidden test data, our CGDE model has achieved higher overall accuracy compared to the NLM-NCBI model (Table 1).

**Table 1.**
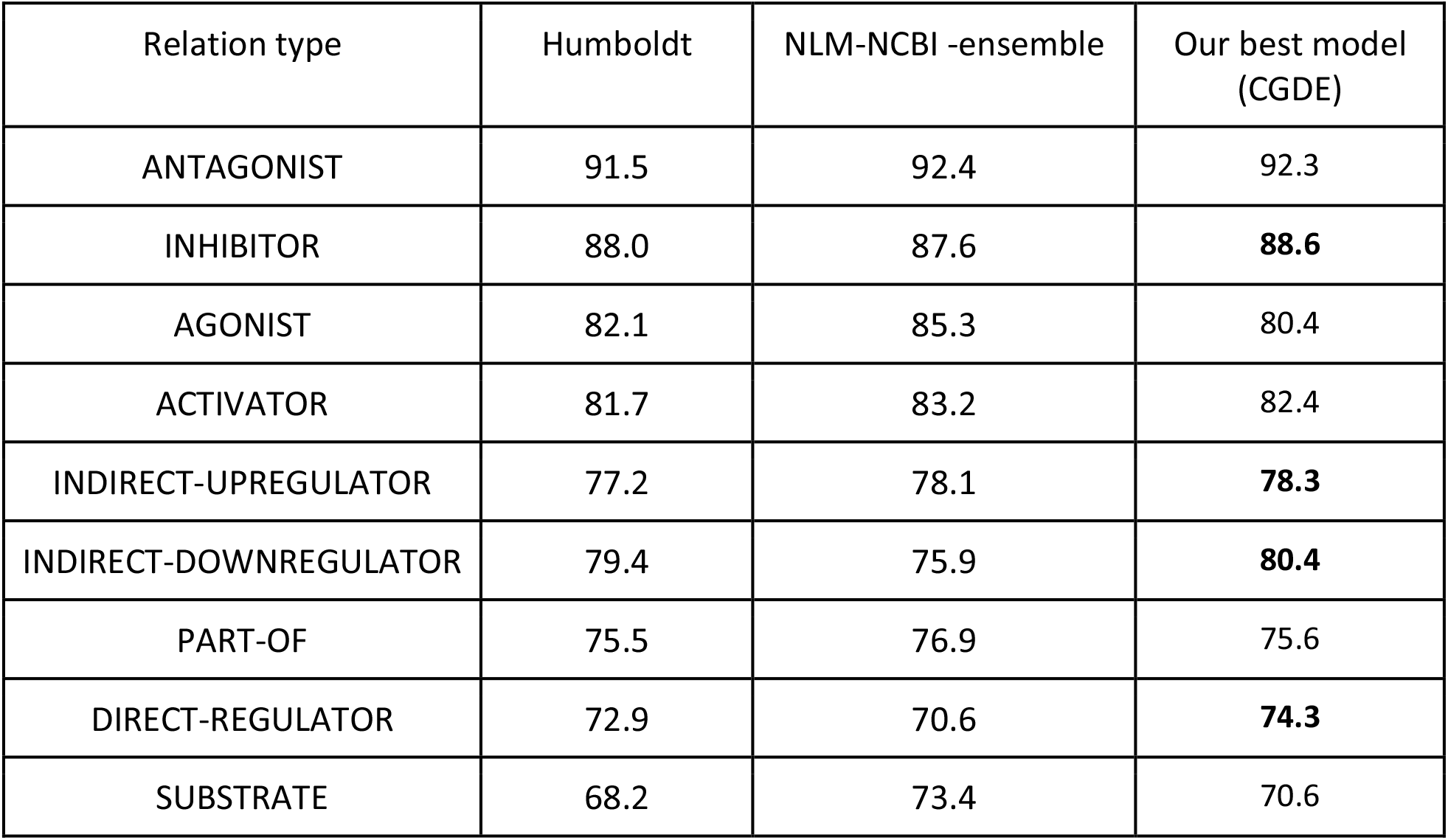
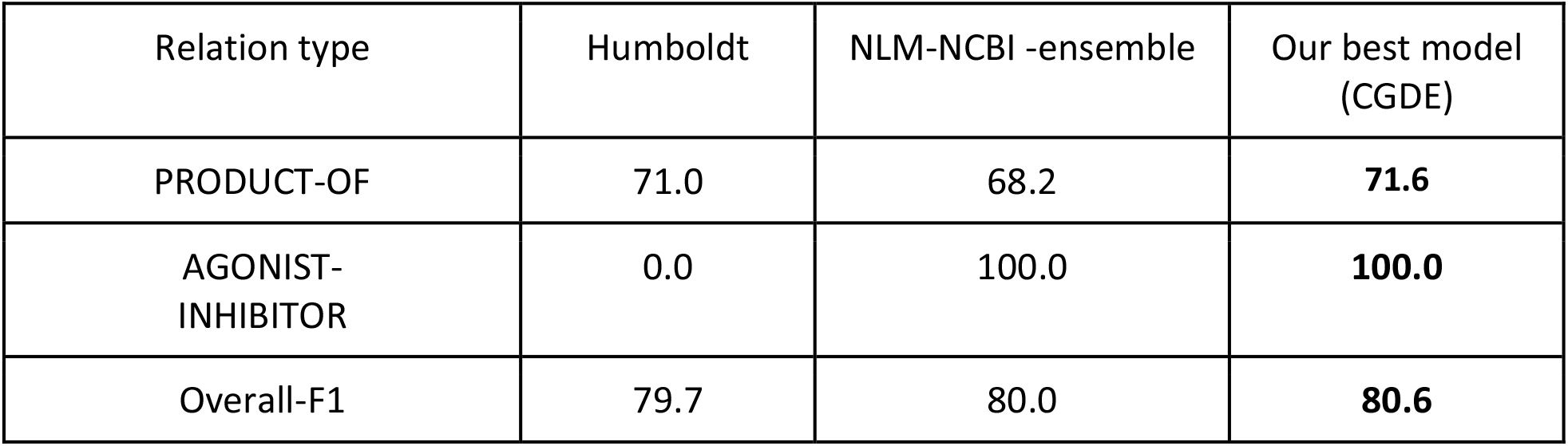
Performance comparison between the NLM-NCBI Ensemble model, and our best model (CGDE) on the DrugProt test data.

| Relation type | Humboldt | NLM-NCBI -ensemble | Our best model (CGDE) |
| --- | --- | --- | --- |
| ANTAGONIST | 91.5 | 92.4 | 92.3 |
| INHIBITOR | 88.0 | 87.6 | **88.6** |
| AGONIST | 82.1 | 85.3 | 80.4 |
| ACTIVATOR | 81.7 | 83.2 | 82.4 |
| INDIRECT-UPREGULATOR | 77.2 | 78.1 | **78.3** |
| INDIRECT-DOWNREGULATOR | 79.4 | 75.9 | **80.4** |
| PART-OF | 75.5 | 76.9 | 75.6 |
| DIRECT-REGULATOR | 72.9 | 70.6 | **74.3** |
| SUBSTRATE | 68.2 | 73.4 | 70.6 |
| PRODUCT-OF | 71.0 | 68.2 | **71.6** |
| AGONIST-INHIBITOR | 0.0 | 100.0 | **100.0** |
| Overall-F1 | 79.7 | 80.0 | **80.6** |

In assessing our CGDE ensemble model, alongside the Humboldt (CDE) ensemble model and the NLM-NCBI ensemble model, we observed a significant variance in the class-wise results.

For the ANTAGONIST class, our model demonstrated high efficacy, with a success rate of 92.3. This was nearly on par with the NLM-NCBI ensemble model at 92.4, but markedly superior to that of the Humboldt CDE model, which scored 91.5. In the INHIBITOR class, our CGDE model outstripped the competition, achieving an 88.6 score. This surpassed the NLM-NCBI model’s 87.6 and the 88 score of the Humboldt CDE model. This pattern was mirrored in the INDIRECT-DOWNREGULATOR, DIRECT-REGULATOR, and PRODUCT-OF relations categories.

However, a different trend emerged for the AGONIST class. Our model lagged slightly behind at 80.4, compared to the NLM-NCBI and Humboldt ensemble models. Similarly, in the ACTIVATOR class, while our model outpaced the Humboldt ensemble model, it was slightly behind the NLM-NCBI ensemble model at 82.4. In the INDIRECT-UPREGULATOR class, our model performed comparably with the NLM-NCBI model, with a marginal 0.2 F1 points advantage, and surpassed the Humboldt ensemble model. Lastly, in the AGONIST-INHIBITOR class, both models achieved a flawless score of 100.

Despite the class-specific variations, the overall F1-score for our CGDE ensemble model model narrowly topped the list with an overall score of 80.6, compared to the NLM-NCBI model’s 80.0 and Humboldt CDE model’s 79.7.

### Comparative analysis: gene-based descriptor vs. sentence-based models

In our study, we investigated the influence of entity descriptions on predictive performance by comparing two distinct models: the sentence-based ensemble model using sentences only and an extended model incorporating gene descriptions. Our analysis conducted on the development dataset yielded valuable insights into their performance. The results of both ensemble models are provided in **Supplementary File 1**. As shown in Figure 6, in terms of True Positives (TPs), we observed that the extended model with gene descriptions outperformed the sentence-based model, with **2966** TPs compared to **2367** TPs. This suggests that gene descriptions provide a substantial benefit in enhancing the predictive capabilities of the model. However, it’s important to note that the extended model also exhibited a higher number of False Positives (FPs) at **642**, as opposed to the baseline model’s **245** FPs. This highlights a trade-off between sensitivity and specificity when incorporating gene descriptions.

**Figure 6.**
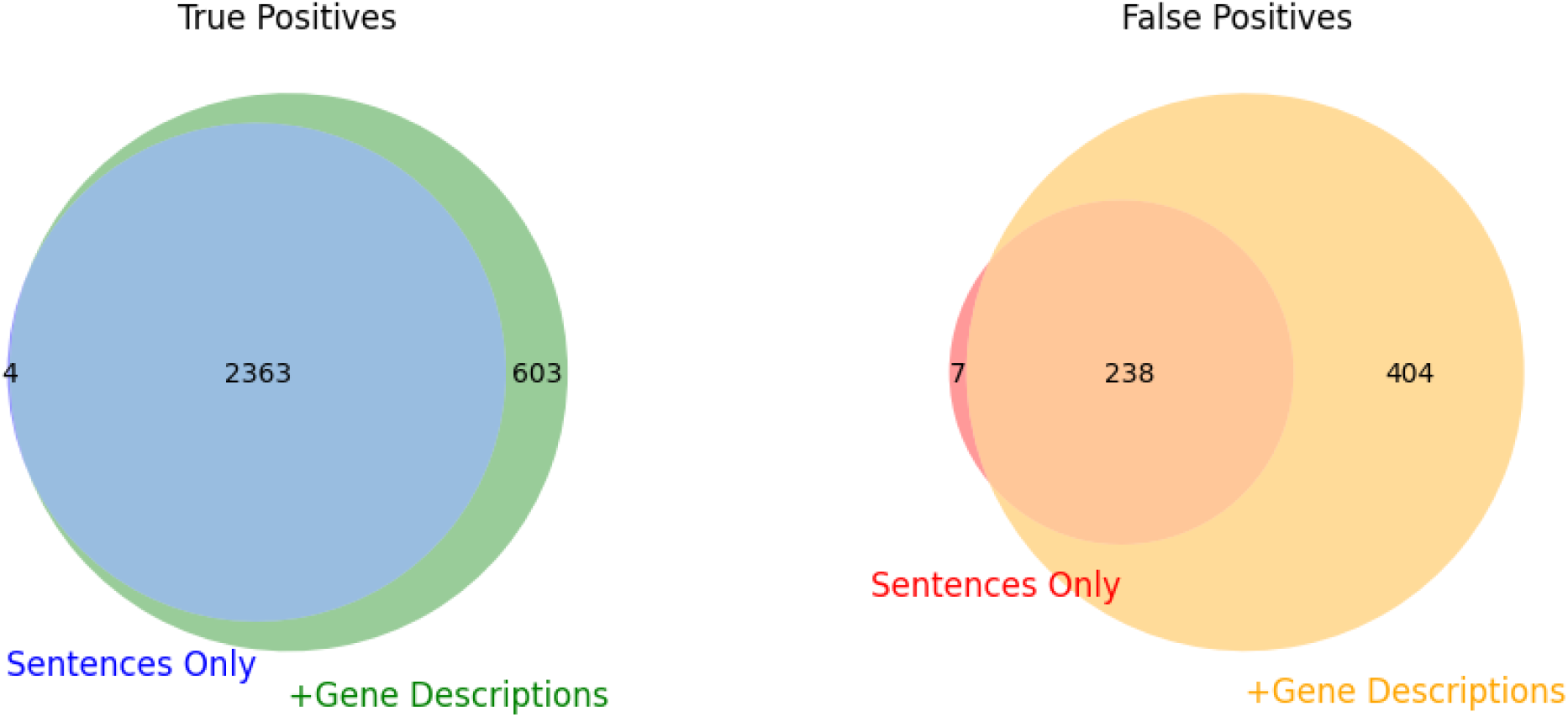
Comparative Analysis of True and False Positives in sentence-based vs. enhanced gene description models.

The overlap in TPs between both models was substantial, with **2363** TPs in common. Notably, the sentence-based ensemble model identified four unique TPs that the extended model did not, while the extended model with gene descriptions identified **603** unique TPs not found by the baseline model. These unique contributions emphasize the added value of including gene descriptions in the model.

We also employed a comprehensive relation type analysis and error distribution analysis. The examination of TPs revealed that the extended model, enriched with gene descriptions, demonstrated substantial improvements across various relation types. Specifically, it surpassed the baseline model in identifying ACTIVATOR and AGONIST relations with **183** TPs and **95** TPs, respectively, compared to the baseline’s **141** and **76**. While both models performed comparably in ANTAGONIST relations, the extended model excelled in identifying DIRECT-REGULATOR relations with an impressive **280** TPs compared to the baseline’s **131**. Additionally, the extended model outperformed the baseline in detecting INDIRECT-DOWNREGULATOR relations, tallying **265** TPs to the baseline’s **188**. However, this enhancement in True Positives came at the cost of increased False Positives (FPs). The extended model exhibited more FPs in ACTIVATOR, AGONIST, ANTAGONIST, DIRECT-REGULATOR, and INDIRECT-DOWNREGULATOR relations, highlighting a potential trade-off between sensitivity and specificity. Notably, the significant improvement in identifying DIRECT-REGULATOR relations by the extended model underscores the added value of gene descriptions in this specific category. Despite the rise in FPs, our analysis suggests that gene description information proves beneficial, particularly for specific relation types, offering valuable insights into the influence of entity descriptions on our predictive models.

### Case Study Analysis: Ozone and IL-6 Interaction

In this case study, we delve into the interaction between Ozone and the IL-6 gene, highlighting the differential predictive capabilities of the sentence-based and gene-based descriptor models. The sentence from the dataset that we focus on is shown in Figure 7. This sentence mentions the effect of Ozone exposure on various genes, including IL-6, but does not explicitly state the nature of the interaction.

**Figure 7.**
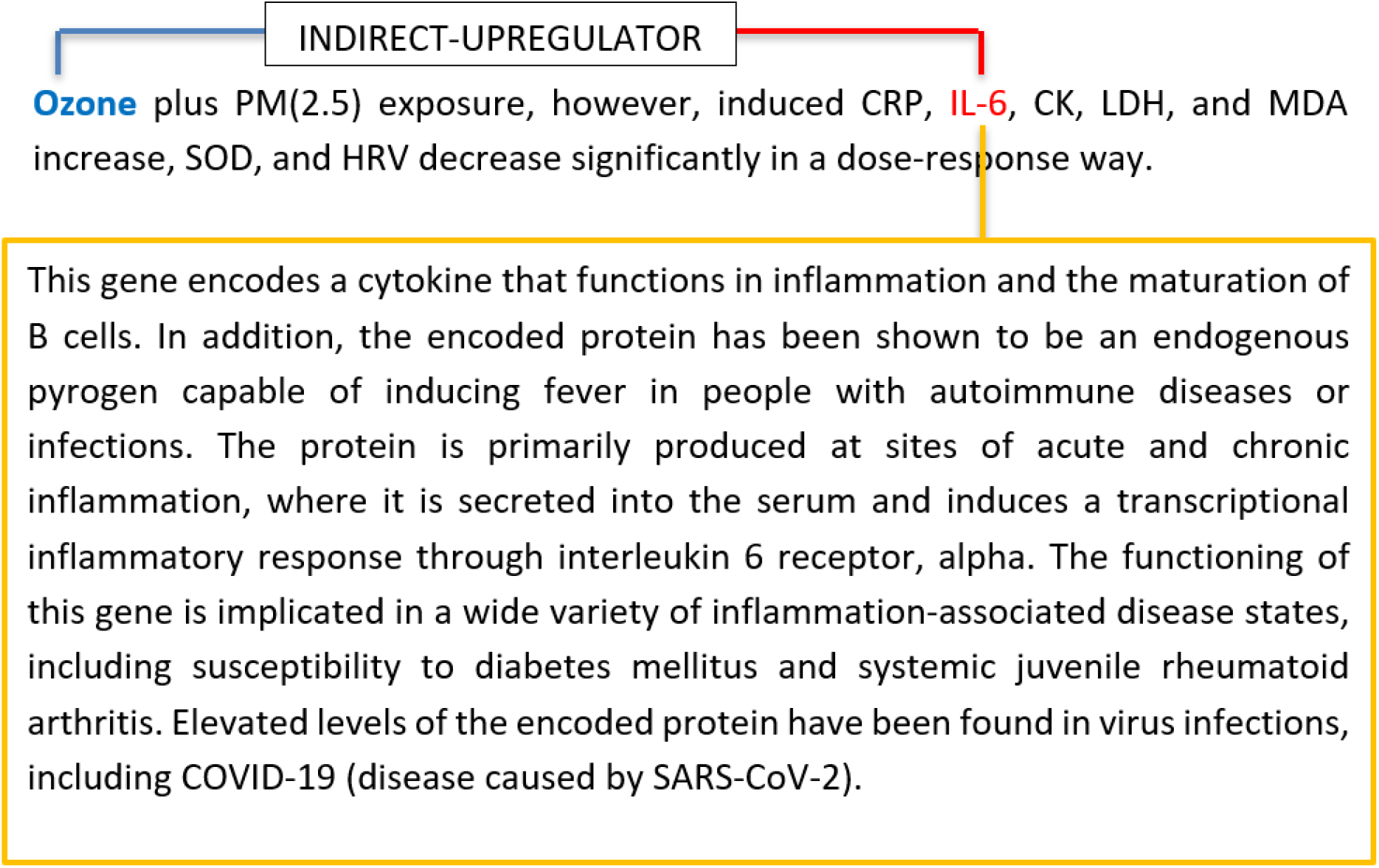
A case study analysis: Ozone and IL-6 interaction. Gene summary description extracted from NCBI Entrez database are highlighted in the rectangle.

We found that sentence-Based Model, relying solely on the sentence, was unable to predict the relationship between Ozone and IL-6 due to the lack of explicit information. In contrast,The Gene-Based Descriptor Model whichincorporates the gene description information, correctly predicted the relationship as an INDIRECT-UPREGULATOR. It inferred this interaction by understanding the role of IL-6 in inflammation and its potential upregulation in response to environmental stressors like Ozone.

### Comparison of NCBI Entrez gene summary and UniProt protein function description

The UniProt protein function is a feature available in the UniProt database [10], a comprehensive resource for protein sequence and functional information. The protein function section provides a brief and informative summary of the protein’s function including the protein’s biological activity, localization, interaction partners, and related pathways. An example of the data is shown in Figure 2B.

The Humboldt team utilized multiple single models using RoBERTa-large-PM-M3-Voc with different seeds, in conjunction with an ensemble model that integrated UniProt protein function descriptions. Their testing ground was the DrugProt development dataset and reported their single results as the mean and standard deviation of the top three runs from a pool of five different random seeds. For our part, we employed a similar methodology, using a variety of single models using RoBERTa-large-PM-M3-Voc and an ensemble model. However, our findings indicated that the NCBI Entrez gene summary data offered a more comprehensive resource, and consequently resulted in slightly better F1 scores when contrasted with the UniProt protein function (Table 2). The results demonstrate that a single model employing gene summary description attains a superior F1 score, showing an improvement of +0.7 points, in contrast to the performance of three model runs that utilized UniProt protein function.

**Table 2.** Comparative analysis of including the UniProt protein function summaries and the NCBI Entrez Gene summary data.

|  | Single |  |  | Ensemble |  |  |
| --- | --- | --- | --- | --- | --- | --- |
|  | P | R | F1 | P | R | F1 |
| **UniProt protein function**<br>**3 runs** | 77.4 ±0.6 | 79± 0.5 | 78.5 ±0.2 | 78.8 | 79.6 | **79.2** |
| **NCBI Entrez gene summary**<br>**1 run** | 79.8 | 79 | **79.4** | - | - | - |
| **NCBI Entrez gene summary**<br>**3 runs** | 79.9 ±1.6 | 77.9 ±1.7 | **78.9 ±0.4** | 79.6 | 79.5 | **79.5** |

## Conclusions

Data deposition from drug target studies into a public database, such as ChEMBL [20], PubChem [21], and Drug Target Commons (DTC) [22], still requires manual curation which is labor-intensive and time-consuming. To keep pace with the rapid growth of the literature, automated methods for extracting DTIs are essential for reducing the time and cost associated with drug discovery and development. However, predicting the drug-target interaction types from scientific literature remains a demanding task. Our study demonstrates the efficacy of incorporating gene and chemical descriptions for the extraction of drug-target interactions from biomedical literature. The proposed methodology significantly improves upon the previous state-of-the-art approaches, showcasing the utility of our approach in biomedical text mining and in enhancing drug discovery efforts. Furthermore, the comparison between the usage of the NCBI Entrez gene summary and the UniProt protein function description indicates that the gene summary offers more comprehensive information about the drug-target interactions, yielding better model performance.

Several opportunities remain for further enhancements. For instance, our experiments have shown that the inclusion of gene sequence-based features did not yield significant improvement in the performance. However, future studies could replace sequence-based features with transcriptomic features as adapted in [23]. Furthermore, we are planning to mine drug-target entities from the 0.3M articles identified in our previous work in [3]. In addition to that, while our method achieved the best performance in the BioCreative DrugProt task, we believe there is still room for improvement in the model architecture. For example, we intend to investigate the potential of leveraging other transformer-based models like BioGPT [24].

Lastly, the developed approach and the conclusions drawn are specific to the extraction of drug-target interactions. It would be interesting to investigate the generalizability of this method for other types of relationship extraction applications in drug discovery such as protein-protein interaction and drug-drug interactions. The results from this study thus represent a significant stride in the field of biomedical text mining and drug discovery, and we believe that the continuous advancements in natural language processing hold immense promise for further progress.

## Key Points

- We leveraged pre-trained NLP models like RoBERTa-large-PM-M3-Voc to efficiently mine drug-target interactions (DTIs) from biomedical texts, significantly advancing drug discovery efforts.
- Our ensemble approach (CGDE), integrating gene and chemical descriptions from Entrez Gene and CTD databases, achieved a leading F1 score of 80.6, demonstrating its effectiveness in DTI extraction.
- We conducted a comprehensive comparative analysis, evaluating the effectiveness of gene textual descriptions sourced from both the Entrez Gene and UniProt databases, to discern their impact on model performance in DTI tasks.
- By incorporating gene descriptions, we improved predictive capabilities, despite a trade-off between sensitivity and specificity, underscoring the value of detailed gene information in DTI extraction.

### Jehad Aldahdooh

is a PhD student at University of Helsinki. He is developing text mining applications for drug–target interactions.

### Ziaurrehman Tanoli

is senior researcher at University of Helsinki. His research is mostly focused on computational drug repurposing. He is also developing bioinformatics tools for drug–target interactions.

### Jing Tang

is an assistant professor at University of Helsinki. He is working on mathematical, statistical and informatics tools to tackle biomedical questions.

## Supporting information

Performance of the SBE and GDE models on the DrugProt development data

