## Supplementary material for "Mining drug-target interactions from biomedical literature using chemical and gene descriptions-based ensemble transformer model": Performance of the SBE and GDE models on the DrugProt development data

The comparison between the Gene Description Ensemble (GDE) and the Sentence-Based Ensemble (SBE) models offers valuable insights into their performance variations on the DrugProt development relation prediction task. The GDE Model, which utilizes gene descriptions in conjunction with an ensemble model consisting of five various RoBERTa-large-PM-M3-Voc models, achieved an impressive global F1 score of 80.6 (Figure 1). This highlights its ability to incorporate biological context effectively, resulting in a well-balanced precision and recall across various relation types. Notably, the GDE model excelled in identifying INHIBITOR and ANTAGONIST relations as shown in Figure 2, achieving an F1 score of 91.3 and 87.7, showcasing its proficiency in accurately recognizing these interactions.

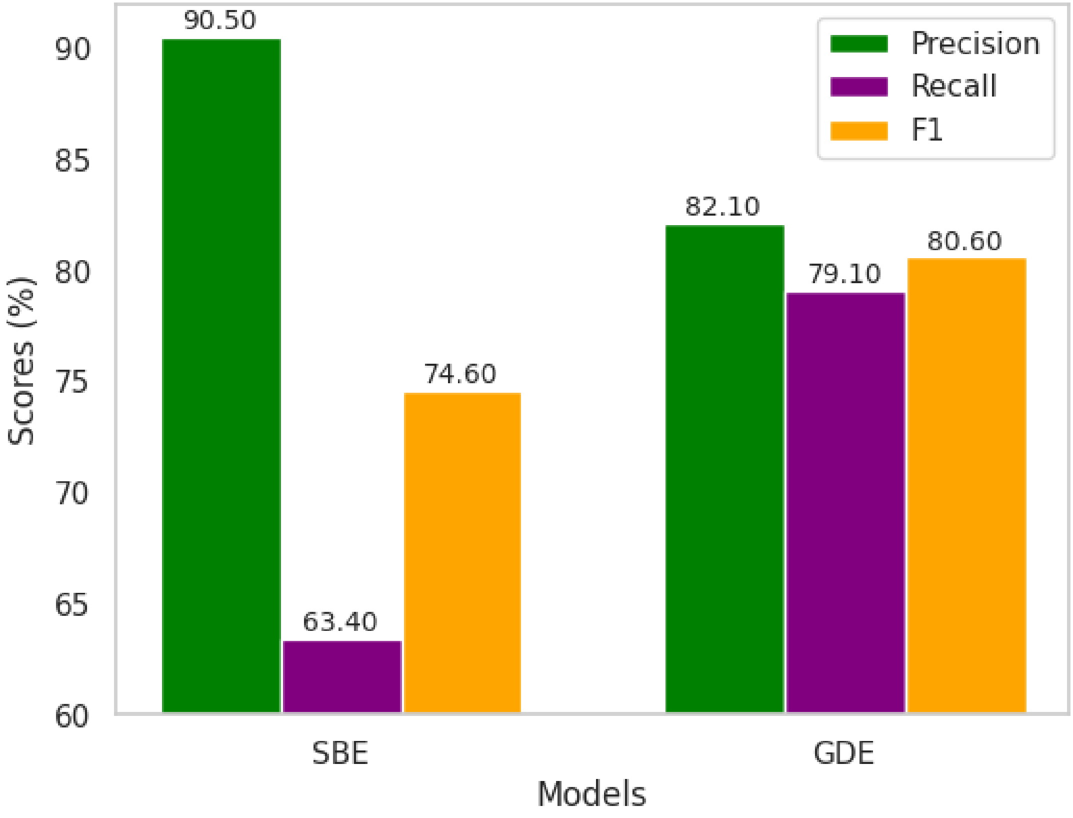


Figure 1: Performance comparison of the SBE and GDE models on the DrugProt development set, presented as micro-averaged precision, recall, and F1 scores, separately.

In contrast, the SBE model, relying solely on sentence-based inputs without external gene or chemical descriptions, exhibited higher precision on a global scale (Figure 1), but faced challenges in terms of recall (63.4), resulting in a lower global F1 score of 74.6 compared to the GDE Model. This discrepancy emphasizes the crucial role of incorporating contextual gene descriptions during model training, as evidenced by the superior performance of the GDE Model. However, it's worth noting that the SBE model demonstrated exceptional precision when predicting the ANTAGONIST relation, achieving the highest F1 score of 90.4 among all the relations it predicted. This suggests that for specific interaction types, the sentence-based approach can be highly effective, especially when precision takes precedence over recall.

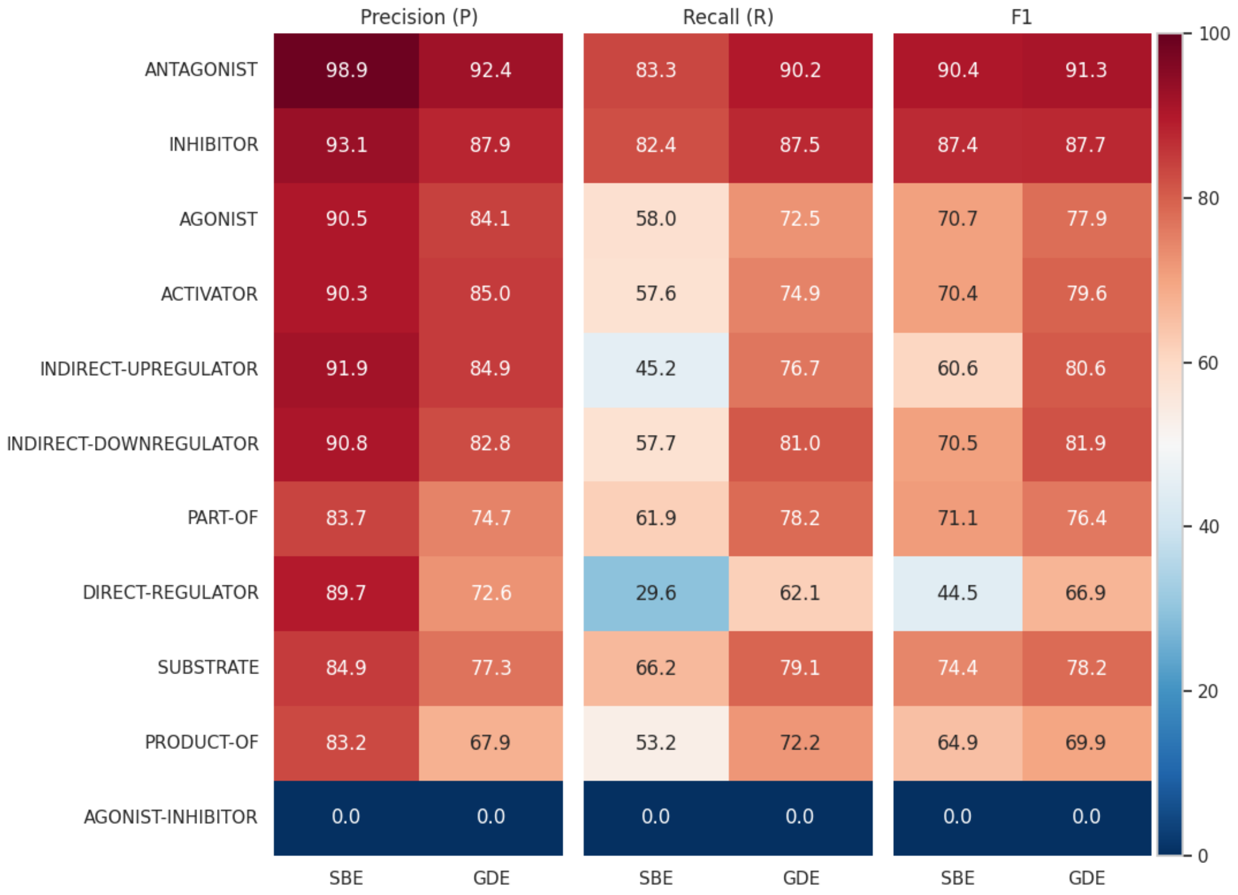


Figure 2: Comparative analysis of model performance per relation types.

A noteworthy observation is the challenge posed by rare classes such as AGONIST-ACTIVATOR, SUBSTRATE_PRODUCT-OF and AGONIST-INHIBITOR relations, where both models reported zero scores. This underscores a common issue in predictive modeling, where limited training samples for specific classes can significantly impact model performance. This highlights the pressing need for more robust training datasets that encompass a broader range of interaction types to enhance the generalizability of these models.

The evident trade-off between precision and recall observed in both models highlights the inherent complexity of balancing these metrics in predictive modeling. For example, while the SBE model leans towards higher precision, it does so at the expense of recall, particularly in relations like DIRECT-REGULATOR, where it achieved a high precision of 89.7 but a low recall of 29.6. This indicates a conservative prediction strategy that minimizes false positives but may miss true positives. In contrast, the GDE model demonstrates a more balanced approach, evident in its overall performance metrics, suggesting that incorporating gene descriptions provides a more comprehensive understanding of interactions, leading to improved recall without significantly compromising precision.
